# Plant-associated *Pseudomonas aeruginosa* harbor multiple virulence traits essential for mammalian infection

**DOI:** 10.1101/2021.07.20.453120

**Authors:** Sakthivel Ambreetha, Ponnusamy Marimuthu, Kalai Mathee, Dananjeyan Balachandar

## Abstract

*Pseudomonas aeruginosa* is a leading opportunistic pathogen capable of causing fatal infections in immunocompromised individuals and patients with degenerative lung diseases. Agricultural soil and plants are the vast reservoirs of this dreaded pathogen. However, there have been limited attempts to analyze the pathogenicity of *P. aeruginosa* strains associated with edible vegetable plants. This study aims to elucidate the virulence attributes of *P. aeruginosa* strains isolated from the rhizosphere and endophytic niches of cucumber, tomato, eggplant, and chili collected from agricultural fields. Virulence of the agricultural strains was compared to three previously characterized clinical isolates. Our results showed that 50% of the plant-associated strains formed significant levels of biofilm and exhibited swarming motility. Nearly 80% of these strains produced considerable levels of rhamnolipid and exhibited at least one type of lytic activity (hemolysis, proteolysis, and lipolysis). Their virulence was also assessed based on their ability to suppress the growth of plant pathogens (*Xanthomonas oryzae, Pythium aphanidermatum, Rhizoctonia solani*, and *Fusarium oxysporum*) and kill a select nematode (*Caenorhabditis elegans*). The plant-associated strains showed significantly higher virulence against the bacterial phytopathogen whereas the clinical strains had significantly higher antagonism against the fungal pathogens. In *C. elegans* slow-killing assay, the clinical strains caused 50-100% death while a maximum of 40% mortality was induced by the agricultural strains. This study demonstrates that some of the *P. aeruginosa* strains associated with edible plants harbor multiple virulence traits. Upon infection of humans or animals, these strains may evolve to be more pathogenic and pose a significant health hazard.

## Introduction

*Pseudomonas aeruginosa* is a leading opportunistic pathogen that causes hospital-acquired, often fatal infections in immunocompromised individuals and patients with chronic pulmonary conditions (Reynolds et al., 1975; Von Graevenitz, 1977; Rosenthal et al., 2020). Additionally, this pathogen can manifest as a wide variety of infections, such as folliculitis, endocarditis, osteomyelitis, and sclerokeratitis, in healthy individuals (Radford et al., 2000; Tate et al., 2003; Doustdar et al., 2019). *P. aeruginosa*-associated mortality is a global concern in healthcare settings, which is why this bacterium is listed among the ‘serious threat pathogens’ (CDC AR, 2019; WHO News, 2019; PHE, 2020).

*P. aeruginosa* is a well-known soil bacterium predominantly found in agricultural ecosystems (Clara, 1930; Elrod and Braun, 1942; Ali Siddiqui and Ehteshamul-Haque, 2001; Adesemoye and Ugoji, 2009; Mondal et al., 2012; Gao et al., 2014; Yasmin et al., 2014; Radhapriya et al., 2015; Arif et al., 2016; Durairaj et al., 2017; Tiwari and Singh, 2017; Gupta and Buch, 2019; Chandra et al., 2020). Few studies have argued that soil and plants are the primary sources for transmission of *P. aeruginosa* to humans (Green et al., 1974; Cho et al., 1975). Plant-associated *P. aeruginosa* first became a significant concern when its presence was detected in fresh vegetables in hospital kitchens, canteens, agricultural farms, retail markets, and supermarkets (Kominos et al., 1972; Wright et al., 1976; Correa et al., 1991; Viswanathan and Kaur, 2001; Curran et al., 2005; Allydice-Francis and Brown, 2012; Ambreetha et al., 2021).

To date, very limited studies have demonstrated the inter-kingdom pathogenicity of *P. aeruginosa* strains. A clinical strain*, P. aeruginosa* PA14 isolated from a hospital burn ward (Mathee, 2018), was reported to elicit extensive rotting in vegetable plants, such as cucumber, lettuce, potato, and tomato (Schroth et al., 1977; Schroth et al., 2018), and *Arabidopsis* (Rahme et al., 2000). *P. aeruginosa* strain BP35, isolated from a black pepper plant, is cytotoxic to mammalian A549 cells (Kumar et al., 2013). Clinical strains of *P. aeruginosa* release a multitude of virulence factors, such as pyocyanin, rhamnolipid, elastases, proteases, lipases, hemolysin, pyochelin, and pyoverdine. These assist the bacterium in establishing lethal infections (Balasubramanian et al., 2012; Moradali et al., 2017). However, there is a clear gap in testing the ability of plant-associated strains to produce the virulence factors required for human infection.

In our previous study, we isolated the plant-associated *P. aeruginosa* strains (PPA01- PPA18) from edible vegetable plants (cucumber, tomato, chili, and eggplant) directly from farms in Southern India (Ambreetha et al., 2021). We reported that those PPA strains were evolutionarily related to the tested clinical isolates (ATCC10145, ATCC9027, and PAO1). Both the agricultural and clinical strains had comparable plant-beneficial traits, such as mineral solubilization, ammonification, extracellular release of indole-3 acetic acid, and siderophore. These results triggered a quest to identify the virulence traits shared between agricultural and clinical *P. aeruginosa* strains. In our current study, we have tested the ability of the plant-associated strains to (1) release the virulence factors critical for human infection and (2) cause mortality in microbial systems and the animal model *Caenorhabditis elegans*.

## Results

### Biofilm formation

In clinical settings, biofilm-forming *P. aeruginosa* causes chronic pulmonary infections (Römling et al., 1994; Bjarnsholt et al., 2009). We hypothesized that *P. aeruginosa* strains in agricultural ecosystems can form biofilms. To test this hypothesis, biofilm formation by the *P. aeruginosa* PPA strains was estimated at three time points (24, 48, and 72 h) using crystal violet-microtiter assay (O’Toole, 2011) and presented as the biofilm to planktonic (B:P) ratio (Fig.1). The significance of the difference among the observed values was assessed using the one-way Analysis of Variance (ANOVA) and Duncan’s Multiple Range Test (DMRT; strains that share the same letters do not differ significantly). ATCC9027, a slow biofilm former, showed low levels at 24h but gradually increased after 48h and 72h of incubation. The well-characterized biofilm-forming strains, ATCC10145 and PAO1, had a high B:P ratio at all three time points. Ten (cucumber: PPA01, PPA02, PPA04; tomato: PPA05, PPA07; eggplant: PPA11, PPA12; chili: PP15, PPA16, PPA18) out of the 18 PPA strains were weak biofilm producers, as evidenced by their B:P ratio of less than one. The cucumber and tomato endophytes, PPA03 and PPA08, produced biofilms comparable to the clinical strains, ATCC9027 and ATCC10145, respectively (indicated by the shared alphabets ‘b’ and ‘c’). Among the plant isolates, the top three strains (PPA03/cucumber, PPA08/tomato, and PPA10/tomato) with a high biofilm population were all endophytes.

**Fig. 1.**
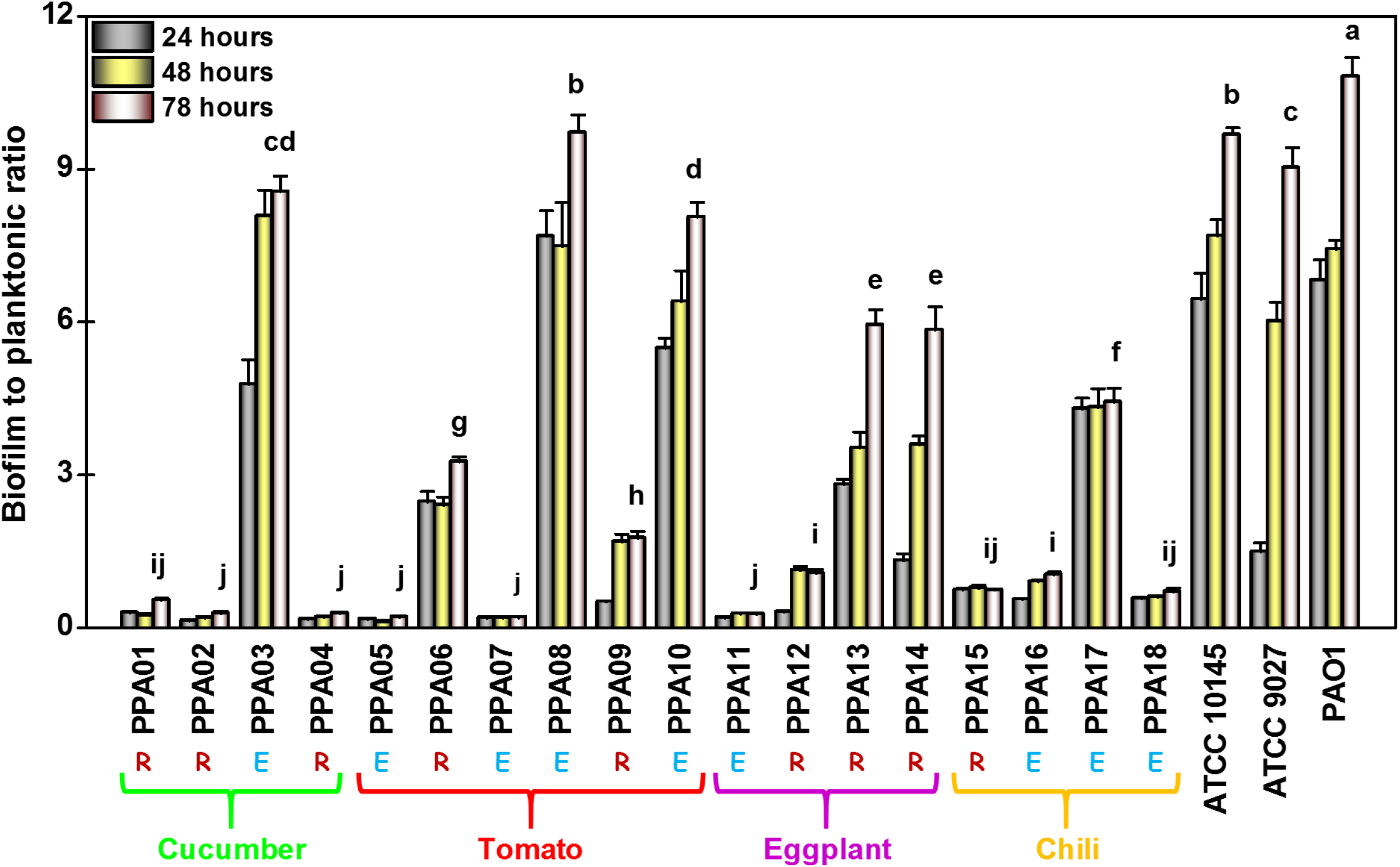
Biofilm production by *P. aeruginosa* strains. The graph represents the biofilm to the planktonic ratio of *P. aeruginosa* strains recorded after 24, 48, and 72 h of incubation. Values plotted are the mean of six replicates with standard errors and letters above the bars indicating the ranking of the strains (significant differences, p < 0.05) based on Duncan’s multiple range test (DMRT). The strains are color-coded based on their plant source: cucumber (green), tomato (red), eggplant (purple), and chili (yellow). The clinical isolates, ATCC10145, ATCC9027, and PAO1 are positive controls. R, rhizosphere strain; E, endophytic strain.

### Swarming motility

Swarming motility is associated with the upregulation of multiple virulence factors in many flagellated bacteria, including *P. aeruginosa* (Overhage et al., 2008; Coleman et al., 2020b; Coleman et al., 2020a). We hypothesized that the plant-associated *P. aeruginosa* strains could exhibit swarming motility. The ability to swarm was assessed using an M9 medium with 0.5% agar (Tremblay and Déziel, 2008). The swarming percentage was calculated based on triple recordings of the diameter of the bacterial tendrils extended on the plate surface. A non-swarming *P. chlororaphis* strain, ZSB15, was used as the negative control (Fig. 2A). The three positive controls, ATCC10145, ATCC9027, and PAO1, spread tendrils that covered more than 50% of the plate’s area (Fig. 2B). All agricultural isolates exhibited swarming patterns at varying levels. The swarming phenotype of the tomato endophyte, PPA08, was significantly higher (covering 80% of the plate) than the tested positive controls, indicated by the letter ‘a’ (Fig.2A and B). Overall, four endophytes (PPA03/cucumber; PPA08, and PPA10/tomato; PPA16, and PPA18/chili), and two rhizospheric strains (PPA13, PPA14/eggplant) were the superior swarmers, swarming more than 50% of plate area. The rest of the strains (PPA01, PPA02, PPA04, and PPA05/cucumber; PPA06, PPAO7, and PPA09/tomato; PPA11, and PPA12/eggplant; PPA15, and PPA17/chili) were weak swarmers. They covered less than 50% of the plate area.

**Fig. 2.**
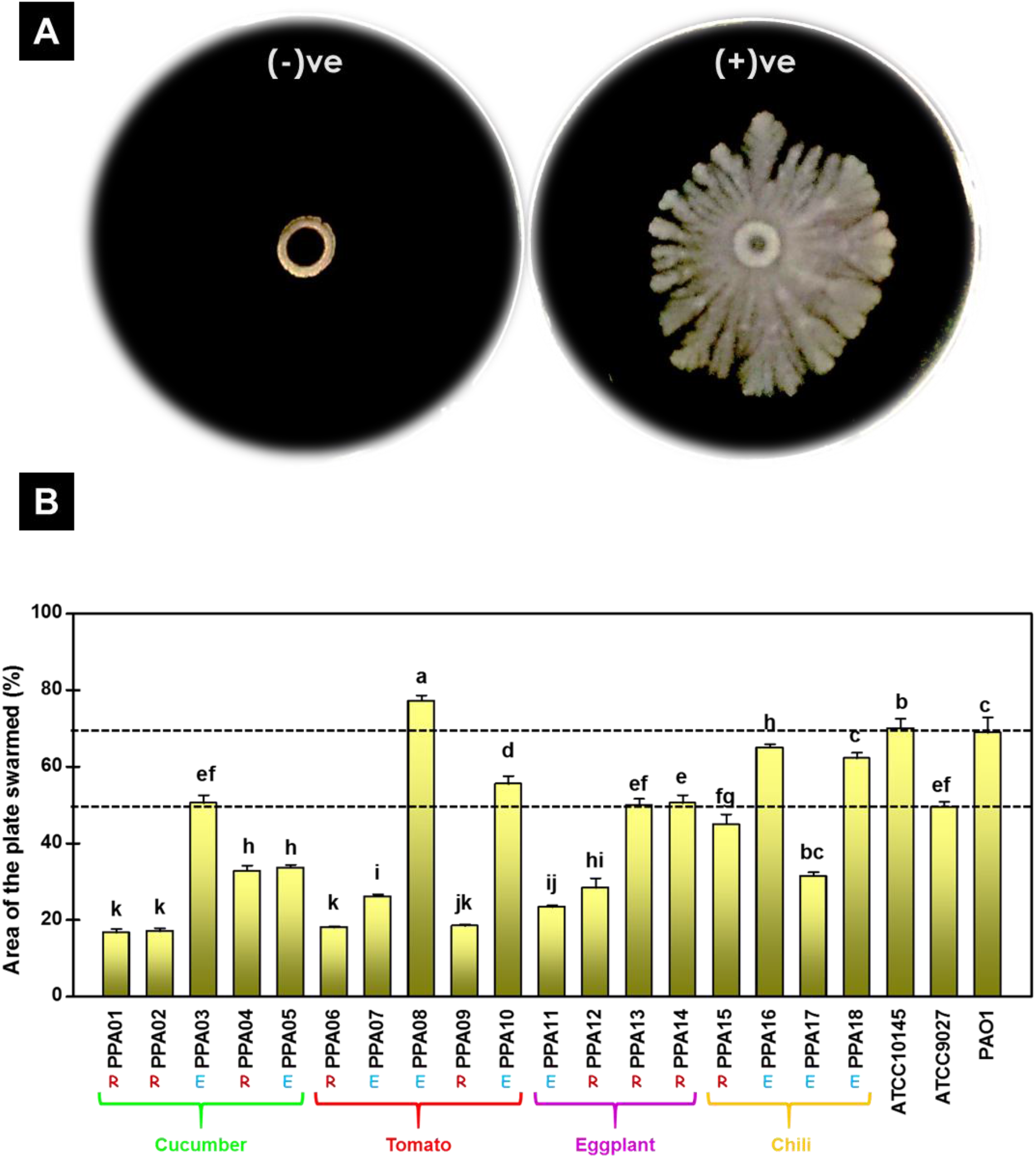
Swarming motility by *P. aeruginosa* strains. (A) Visualization of a non-swarming negative control, *P. chlororaphis* (left), and a superior swarmer, *P. aeruginosa* (PPA08/tomato endophyte), on M9 plates with 0.5% agar. (B) The graph represents the percentage of 90mm Petri-plates covered by the tendrils formed by the *P. aeruginosa* strains during swarming. Values plotted are the mean of six replicates with the standard errors and letters above the bars indicating the ranking of strains (significant differences (p < 0.05) based on Duncan’s multiple range test (DMRT). The strains are color-coded based on their plant source: cucumber (green), tomato (red), eggplant (purple), and chili (yellow). The clinical isolates, ATCC10145, ATCC9027, and PAO1, are positive controls. R, rhizosphere strain; E, endophytic strain.

### Extracellular release of rhamnolipid

Rhamnolipids are a class of metabolites predominantly released by *P. aeruginosa* to infiltrate mammalian lung tissues (McClure and Schiller, 1992, 1996; Zulianello et al., 2006). In plants, rhamnolipids provide protection against pests and pathogens (Kim et al., 2011; Yan et al., 2015; Sancheti and Ju, 2019). In this study, the agricultural strains of *P. aeruginosa* were hypothesized to produce extracellular rhamnolipids. The test strains were qualitatively screened for their ability to release rhamnolipids on cetyltrimethylammonium bromide (CTAB) agar plates (Fig. 3A). All *P. aeruginosa* strains formed blue halo zones around the wells, thus testing positive for rhamnolipid production (Fig. 3A).

**Fig. 3.**
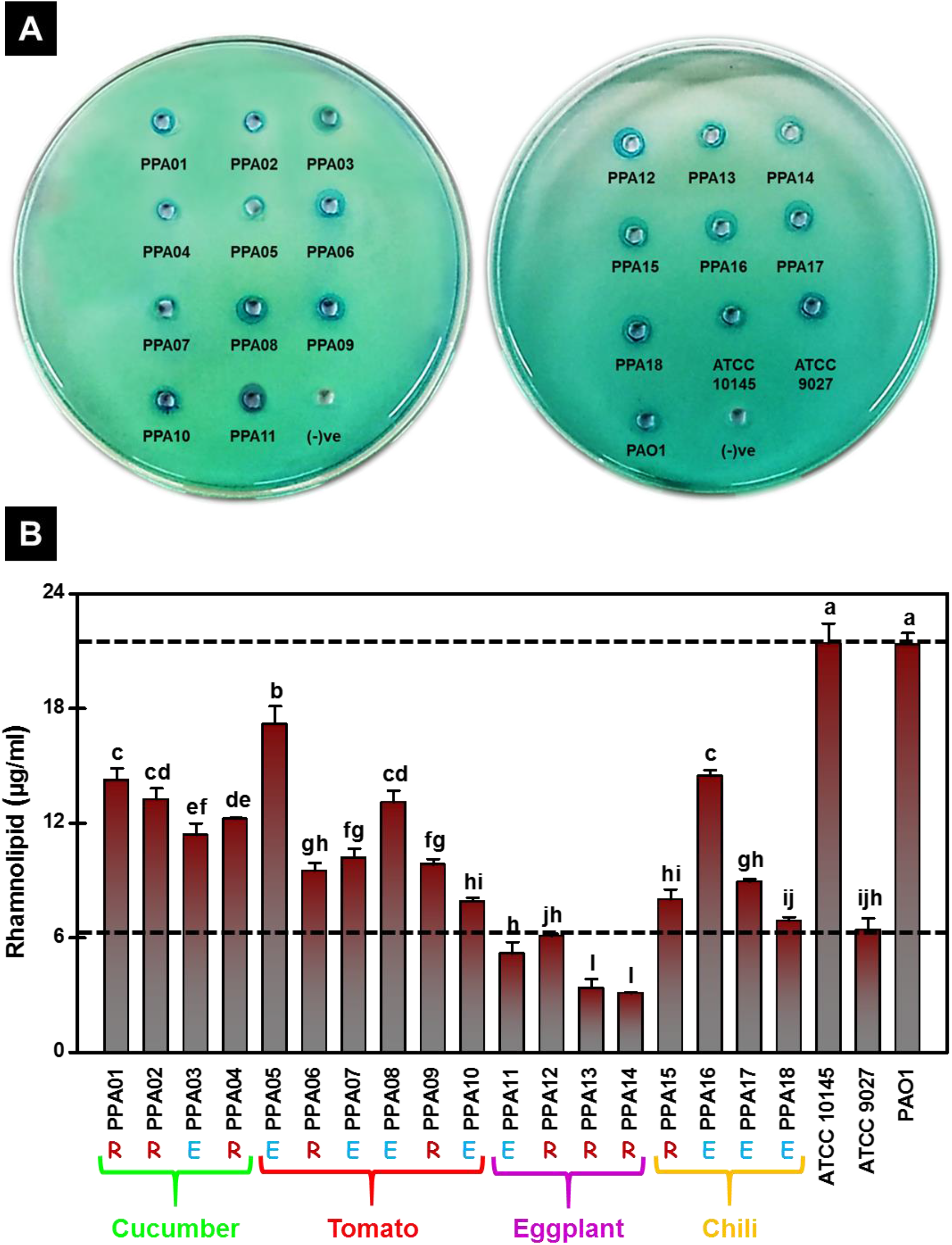
Rhamnolipid production by *P. aeruginosa* strains. (A) Rhamnolipid production is indicated by the appearance of blue halos around the wells upon addition of cell-free supernatant of *P. aeruginosa* strains on CTAB-methylene blue agar medium. (B) Quantitative rhamnolipid levels released by *P. aeruginosa* strains. Values plotted are the mean of three replicates with the standard errors and letters above the bars indicating the ranking of the strains (significant differences (p < 0.05) based on Duncan’s multiple range test (DMRT). The strains are color-coded based on their plant source: cucumber (green), tomato (red), eggplant (purple), and chili (yellow). The dashed lines indicate the levels of rhamnolipid made by the clinical strains, ATCC10145, ATCC9027, and PAO1 (positive controls). R, rhizosphere strain; E, endophytic strain.

Quantitative assessment of extracellular rhamnolipids was performed using the gravimetric method (Zhang and Miller, 1992; Gunther et al., 2005). Rhamnolipid levels were expressed as µg/ml and the statistical significance was expressed through DMRT (Fig. 3B). Two of the three clinical strains, PAO1 and ATCC10145, released a high quantity of rhamnolipids. The clinical isolate ATCC9027 from otitis externa (Table 1) produced comparatively low rhamnolipid levels. All eighteen plant-associated strains released extracellular rhamnolipids. All strains except for the eggplant isolates (PPA11- PPA14) produced more rhamnolipids than ATCC 9027 (Fig. 3B).

**Table 1.** Microbial strains used in this study.

| **Microorganism** | **Source** | **Infection/Niche** | **References** |
| --- | --- | --- | --- |
| ***Pseudomonas aeruginosa* (reference strains)** |  |  |  |
| PAO1 | Human | Wound infection | Holloway, (1955) |
| ATCC9027 | Human | Otitis externa | Haynes, (1951) |
| ATCC10145 | Human | Unknown | Picard et al. (1994) |
| **Plant-associated *P. aeruginosa* strains** |  |  |  |
| PPA01 | Cucumber | Rhizosphere | Ambreetha et al. (2021) |
| PPA02 | Cucumber | Rhizosphere | Ambreetha et al. (2021) |
| PPA03 | Cucumber | Endophyte | Ambreetha et al. (2021) |
| PPA04 | Cucumber | Rhizosphere | Ambreetha et al. (2021) |
| PPA05 | Tomato | Endophyte | Ambreetha et al. (2021) |
| PPA06 | Tomato | Rhizosphere | Ambreetha et al. (2021) |
| PPA07 | Tomato | Endophyte | Ambreetha et al. (2021) |
| PPA08 | Tomato | Endophyte | Ambreetha et al. (2021) |
| PPA09 | Tomato | Rhizosphere | Ambreetha et al. (2021) |
| PPA10 | Tomato | Endophyte | Ambreetha et al. (2021) |
| PPA11 | Eggplant | Endophyte | Ambreetha et al. (2021) |
| PPA12 | Eggplant | Rhizosphere | Ambreetha et al. (2021) |
| PPA13 | Eggplant | Rhizosphere | Ambreetha et al. (2021) |
| PPA14 | Eggplant | Rhizosphere | Ambreetha et al. (2021) |
| PPA15 | Chili | Rhizosphere | Ambreetha et al. (2021) |
| PPA16 | Chili | Endophyte | Ambreetha et al. (2021) |
| PPA17 | Chili | Endophyte | Ambreetha et al. (2021) |
| PPA18 | Chili | Endophyte | Ambreetha et al. (2021) |
| **Phytopathogens** |  |  |  |
| *Xanthomonas oryzae* | - | - | Unpublished |
| *Pythium aphanidermatum* | - | - | Unpublished |
| *Rhizoctonia solani* | - | - | Unpublished |
| *Fusarium oxysporum* | - | - | Unpublished |

### Lytic activity

*P. aeruginosa* lytic enzymes deteriorate pulmonic health by causing vascular permeability, and organ damage (Ostroff et al., 1989; Wargo et al., 2011). We hypothesized that *P. aeruginosa* strains associated with agricultural plants harbor lytic activity. To confirm this, the hemolytic, proteolytic, and lipolytic activities of the strains were qualitatively assessed.

#### Hemolysis

We tested the ability of the *P. aeruginosa* strains to lyse blood on sheep blood agar medium (Williams and Harper, 1947). Strains that partially lysed red blood cells and resulted in a green discoloration on the agar were scored positive for α- hemolytic activity (Table 2). Strains that did not exhibit lytic behavior were marked as γ-hemolytic. As expected, the three control strains, ATCC10145, ATCC9027, and PAO1, exhibited α-hemolytic activity. More than 50% of the agricultural isolates exhibited α- hemolysis, including four rhizospheric strains (PPA02 and PPA04/cucumber; PPA13, and PPA14/eggplant) and six endophytes (PPA03/cucumber; PPA07, PPA08, and PPA10/tomato; PPA11/eggplant; PPA16/chili). The remaining isolates did not exhibit any hemolytic activity.

**Table 2.** Lytic behavior of P. aeruginosa strains.

| Strains | Hemolysis |  | Proteolysis |  | Lipolysis |
| --- | --- | --- | --- | --- | --- |
| | γ-hemolysis | α-hemolysis | Casein | Gelatin | Lipid |
| PPA01 | + | - | - | - | - |
| PPA02 | - | ++ | + | - | - |
| PPA03 | - | ++ | ++ | ++ | - |
| PPA04 | - | ++ | + | ++ | - |
| PPA05 | - | - | + | ++ | - |
| PPA06 | + | - | + | - | + |
| PPA07 | - | ++ | ++ | ++ | + |
| PPA08 | - | ++ | ++ | ++ | - |
| PPA09 | + | - | - | - | - |
| PPA10 | - | ++ | ++ | ++ | - |
| PPA11 | - | ++ | + | ++ | - |
| PPA12 | + | - | + | - | - |
| PPA13 | - | ++ | ++ | ++ | + |
| PPA14 | - | ++ | ++ | ++ | - |
| PPA15 | + | - | ++ | ++ | + |
| PPA16 | - | ++ | ++ | ++ | - |
| PPA17 | + | - | ++ | ++ | - |
| PPA18 | + | - | ++ | ++ | - |
| ATCC10145 | - | ++ | ++ | ++ | + |
| ATCC9027 | - | ++ | - | + | - |
| PAO1 | - | ++ | ++ | ++ | ++ |

γ-hemolysis – no lysis of blood cells; α-hemolysis - partial destruction of blood cells; '+'- mild lysis; '++' – extensive lysis; '-' – no lysis

#### Proteolysis

*P. aeruginosa* associated lysis of the proteins casein and gelatin was tested by plate assay (Atlas, 1993; Georgescu et al., 2016) and presented as positive and negative scores (Table 2). Two of the three tested controls (ATCC10145 and PAO1) harbored high proteolytic activity. ATCC9027 caused mild lysis of gelatin but no lysis of casein. 16 out of 18 plant-associated strains showed caseinase activity whereas only 13 strains had gelatinase activity. The rhizospheric strains (PPA01/cucumber and PPA09/tomato) were unable to hydrolyze either protein. Three other rhizospheric strains (PPA02/cucumber; PPA06/tomato; PPA12/eggplant) that displayed low caesinase activity did not exhibit gelatinase activity.

#### Lipolysis

The lipid hydrolytic activity of the *P. aeruginosa* strains was tested in tributyrin agar medium (Atlas, 1993; Georgescu et al., 2016). Two of three control strains, ATCC10145 and PAO1, showed lipolytic behavior while ATCC9027 did not lyse the tested lipid (Table 2). Most plant-associated strains did not exhibit lipolysis except for three rhizospheric strains (PPA06/tomato, PPA13/eggplant, and PPA15/chili) and one endophyte (PPA07/tomato).

### Antagonism against phytopathogens

Agricultural *P. aeruginosa* strains have been previously shown to inhibit other phytopathogens (Ali Siddiqui and Ehteshamul-Haque, 2001; Yasmin et al., 2014; Durairaj et al., 2017). This study hypothesized that both agricultural and clinical *P. aeruginosa* exhibit virulence against plant pathogens. To test this hypothesis, we challenged the *P. aeruginosa* strains with common fungal (*Pythium aphanidermatum*, *Rhizoctonia solani*, and *Fusarium oxysporum*) and bacterial (*Xanthomonas oryzae*) phytopathogens (Sakthivel and Gnanamanickam, 1986). This is the first known attempt to test the antagonism of clinical strains against phytopathogens. Inhibition of phytopathogens caused by the *P. aeruginosa* strains was normalized to PAO1 (Fig. 4).

**Fig 4.**
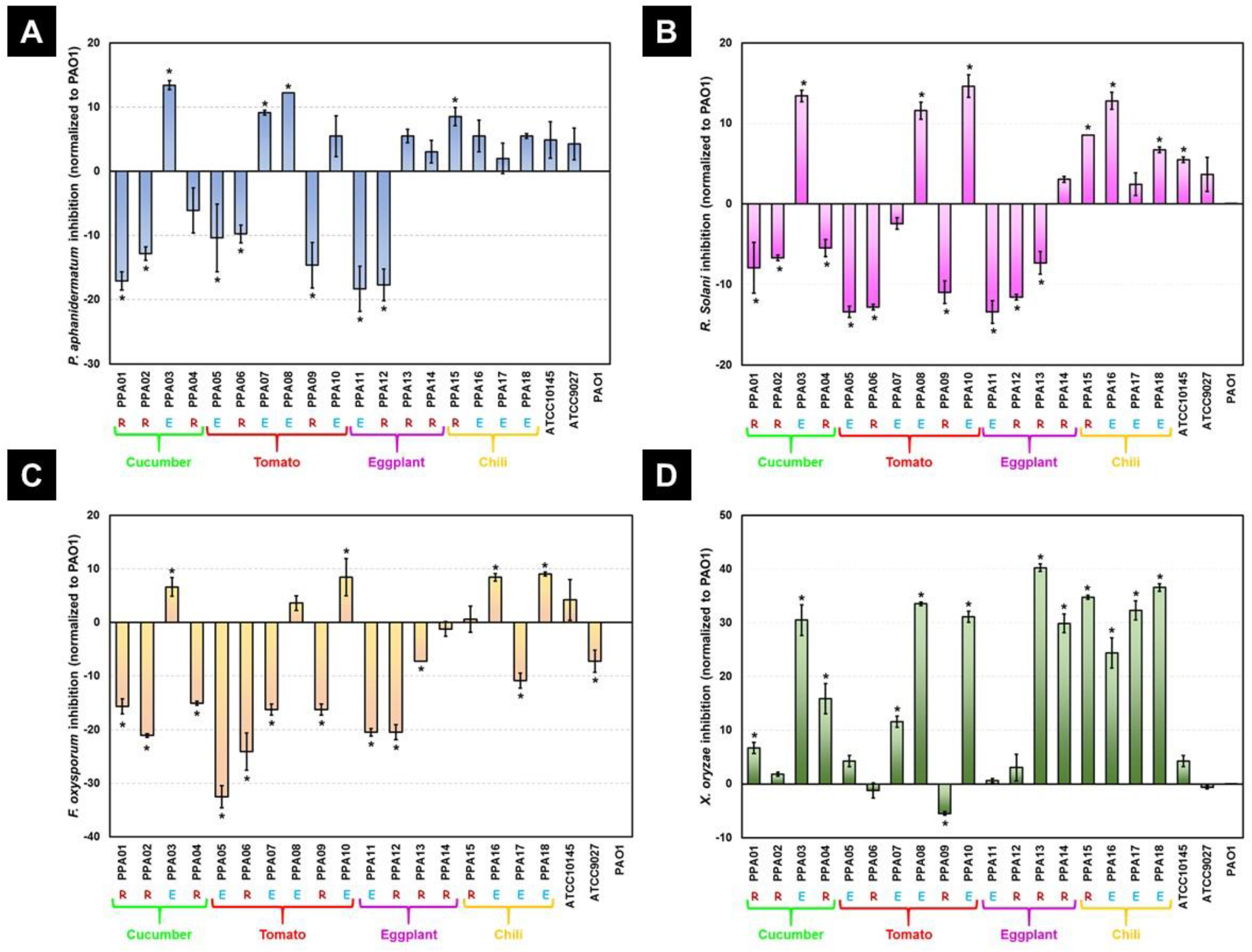
Biocontrol of phytopathogens by *P. aeruginosa* strains. The percentage inhibition of *Pythium aphanidermatum* (A), *Rhizoctonia solani* (B), *Fusarium oxysporum* (C), and *Xanthomonas oryzae* (D) induced by the *P. aeruginosa* strains. Values plotted are the mean of three replicates normalized to PAO1. * denotes the significant difference of PAO1 (p < 0.05) based on Duncan’s multiple range test (DMRT). Strains are color-coded based on their plant source: cucumber (green), tomato (red), eggplant (purple), and chili (yellow). R, rhizosphere strains; E, endophytic strain.

#### Pythium aphanidermatum inhibition

All tested strains could inhibit *Pythium aphanidermatum* (Fig. 4A). Ten plant-associated strains (PPA03/cucumber; PPA07, PAA08, and PPA10/tomato; PPA13, and PPA14/eggplant; PPA15-PPA18/chili) and two clinical strains (ATCC10145 and ATCC9027) showed higher antagonism when compared to PAO1. However, only four PPA strains (PPA03/cucumber; PPA07, and PPA08/tomato; PPA15/chili) were significantly more antagonistic than PAO1 (p<0.05, DMRT). The remaining strains (7 of 18) from cucumber, tomato, and eggplant inhibited *Pythium aphanidermatum* significantly less as compared to PAO1.

#### *R. solani* inhibition

All tested strains could inhibit *R. solani* (Fig. 4B). Eight plant-associated strains (PPA03/cucumber; PAA08, and PPA10/tomato; PPA14/eggplant; PPA15-PPA18/chili) and two clinical strains (ATCC10145 and ATCC9027) showed higher antagonism when compared to PAO1. Among them, six PPA strains (PPA03/cucumber;

PPA08, and PPA10/tomato; PPA15, PPA16, and PPA18/chili) significantly inhibited *R. solani* more than PAO1 (p<0.05, DMRT). The other *P. aeruginosa* strains (9 of 18) from cucumber, tomato, and eggplant caused significantly lower inhibition of *R. solani* than PAO1.

#### F. oxysporum inhibition

All tested strains inhibited *F. oxysporum* (Fig. 4C). Five plant- associated strains (PPA03/cucumber; PAA08, and PPA10/tomato; PPA16, and PPA18/chili) and one clinical strain (ATCC10145) showed higher antagonism when compared to PAO1. Among them, four PPA strains (PPA03/cucumber; PPA10/tomato; PPA16 and PPA18/chili) caused significantly higher inhibition than PAO1 (p<0.05, DMRT). The rest of the *P. aeruginosa* PPA strains (13 out of 18) from cucumber, tomato, eggplant, and chili were significantly less antagonistic against *F. oxysporum* when compared to PAO1.

#### *X. oryze* inhibition

All tested strains could inhibit *X. oryzae* (Fig. 4C). Most of the plant- associated strains (16 out of 18) showed higher antagonism when compared to PAO1. Among them, 12 PPA strains (PPA01, PPA03, and PPA04/cucumber; PPA07, PPA08, and PPA10/tomato; PPA13 and PPA14/eggplant; PPA15-PPA18/chili) caused significantly higher inhibition than PAO1 (p<0.05, DMRT). Two rhizospheric strains (PPA06, and PPA09/tomato) had comparatively lower antagonism of *X. oryzae* than PAO1.

#### Clustering based on antagonistic potential

Euclidean distance-based principal coordinate analysis (PCoA) (NCSS, Kaysville, USA) clustered the *P. aeruginosa* strains based on their combined antagonism against the phytopathogens (Fig. 5). The clinical strains (PAO1, ATCC10145, and ATCC9027) did not cluster with the PPA strains. The PPA strains formed three clusters except for a chili endophyte (PPA16) and tomato rhizospheric strain (PPA09). Cluster A was occupied by three tomato isolates: one from the rhizosphere niche (PPA06) and two endophytes (PPA05 and PPA07). Cluster B contained five endophytes and four rhizosphere strains from all four plants. Cluster C contained four strains isolated from the eggplant and cucumber. Two rhizospheric strains (PPA01 and PPA02) isolated from the cucumber superimposed on each other in cluster C, which reflects their identical antagonism.

**Fig. 5.**
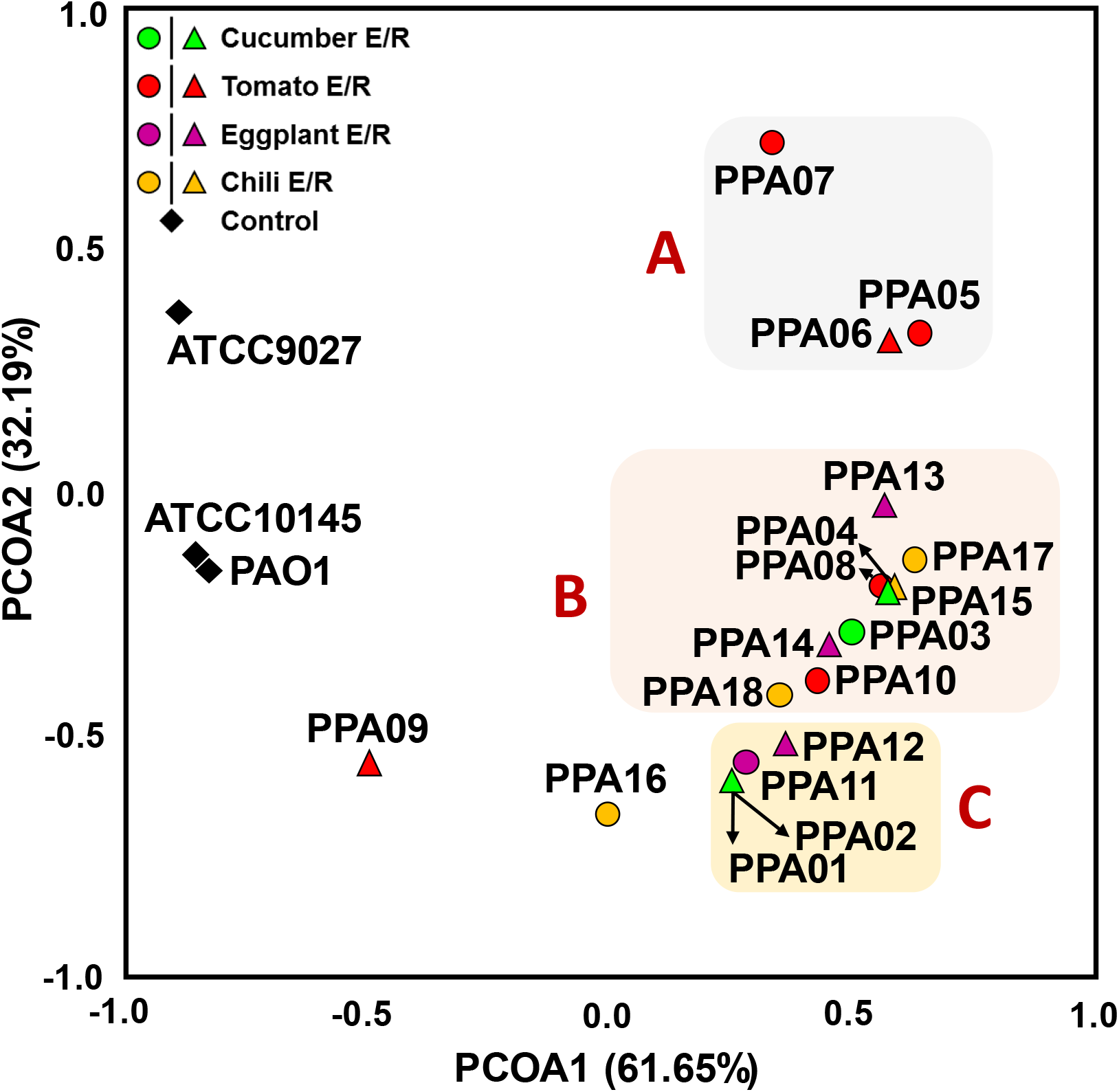
Principal coordinate analysis (PCoA) based on biocontrol ability of the *P. aeruginosa* strains. Euclidean distance-based PCoA plot for the biocontrol of bacterial and fungal phytopathogens by the *P. aeruginosa* strains. The percentage values in parentheses on the x- (PCoA1) and y-axes (PCoA2) depict the similarities and differences among the strains based on their mineral solubilizing ability. The three major clusters of PPA strains formed based on their similar biocontrol activity are named A, B, and C. The strains are color-coded based on their plant source: cucumber (green), tomato (red), eggplant (purple), and chili (yellow).

### Virulence in the animal model, Caenorhabditis elegans

The nematode *C. elegans* has been extensively used as a model system to understand the pathogenicity of *P. aeruginosa* (Mahajan-Miklos et al., 1999; Adonizio et al., 2008). In this study, we hypothesized that the agricultural *P. aeruginosa* strains were capable of killing the *C. elegans* worms. This hypothesis was tested through the *C. elegans* slow killing assay (Tan et al., 1999). The nematodes were scored alive or dead (Fig. 6A) based on their response to physical stimuli. The percentage of living nematodes after feeding on the *P. aeruginosa* strains was noted every 24 h until 120 h of incubation (Fig. 6B). As expected, the negative control, *E. coli* OP50, did not induce mortality in the worms (Fig. 6A). However, at 120 h nearly 8% of the worms died on OP50 plates due to natural death (Fig. 6B). The three positive controls, ATCC10145, ATCC9027, and PAO1, caused higher mortality than the plant-associated strains. All of the nematodes fed with ATCC10145 and PAO1 were dead within 72 and 120 h, respectively. In contrast, only 50% of the worms died after feeding with ATCC9027. Most of the plant-associated strains were less virulent against *C. elegans*. Three endophytic strains from cucumber (PPA03), tomato (PPA08), and chili (PPA18) plants caused maximum mortality of 40%. Only 15% of worms died after feeding on certain rhizospheric (PPA06/tomato; PPA14/eggplant) and endophytic strains (PPA05, and PPA07/tomato). These four strains were the least virulent among the agricultural isolates.

**Fig. 6.**
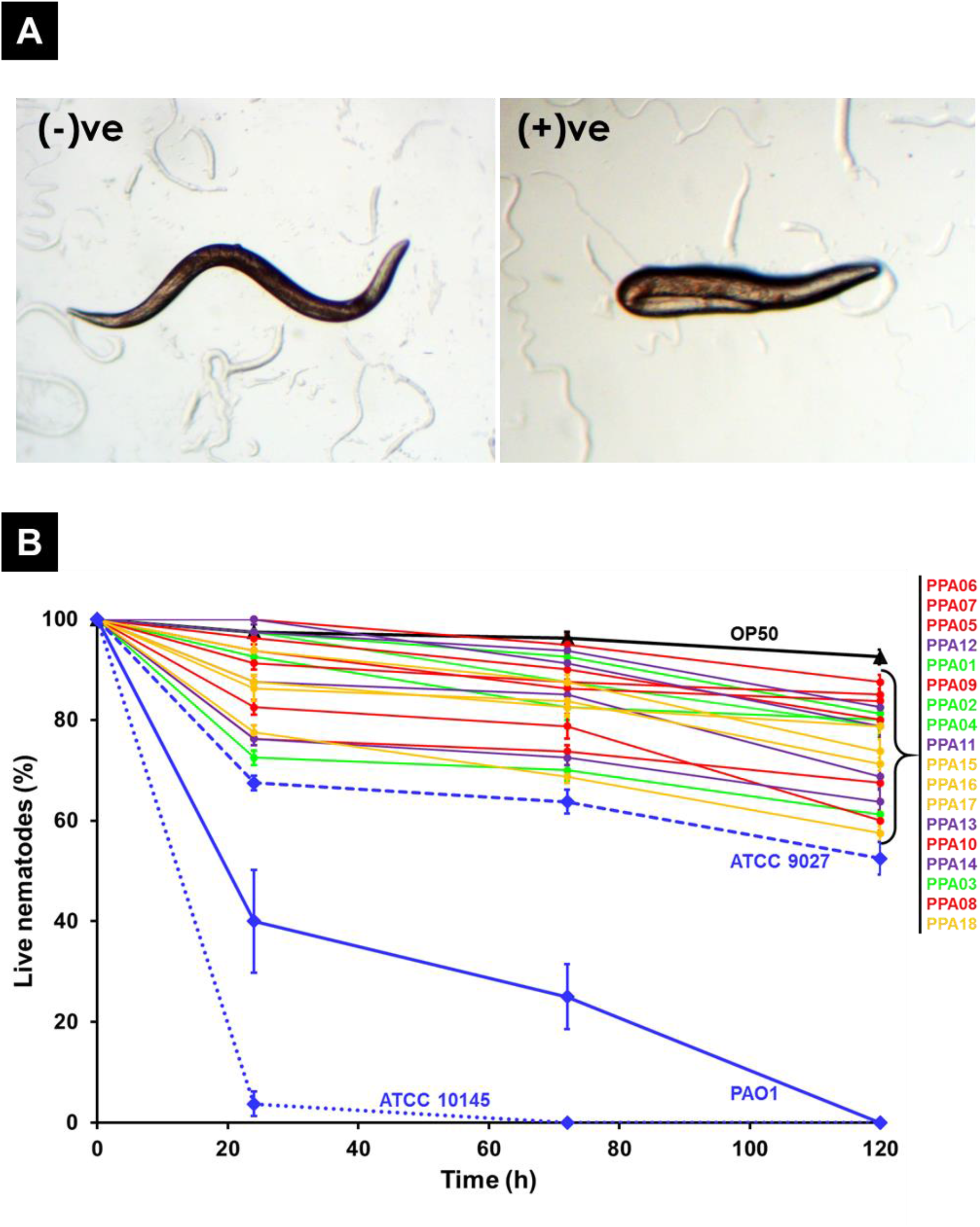
*Caenorhabditis elegans* death induced by *P. aeruginosa* strains. (A) Stereomicroscopic view of L4 nematodes - live and active worm after feeding on *E. coli* OP50 (left); dead worm after feeding on the most virulent clinical isolate, ATCC 10145 (right). (B) Percentage of living nematodes after feeding on *P. aeruginosa* strains recorded over the time course of 0-120 hours. Values plotted are the means of three replicates with standard errors. The blue lines indicate the percentage of nematodes that survived after feeding on the clinical isolates, ATCC10145, ATCC9027, and PAO1 (positive controls). The black line indicates the percentage of living nematodes after feeding on *E. coli* OP50 (negative control). The PPA strains are color-coded based on their plant source: cucumber (green), tomato (red), eggplant (purple), and chili (yellow).

## Discussion

In the 1970s, agricultural soil and plants were recognized as reservoirs of the opportunistic pathogen *P. aeruginosa* (Green et al., 1974; Cho et al., 1975). Since, *P. aeruginosa* has been detected in fresh agricultural produce at markets, hospital kitchens, and local vendors (Kominos et al., 1972; Wright et al., 1976; Correa et al., 1991; Viswanathan and Kaur, 2001; Allydice-Francis and Brown, 2012; Nithya and Babu, 2017). Despite these reports, there have been minimal attempts to characterize the pathogenicity of the plant-associated *P. aeruginosa* strains (Lebeda et al., 1984; Kumar et al., 2013). Our previous study demonstrated that *P. aeruginosa* strains (PPA01 to PPA18) present in the endophytic and rhizospheric niches of cucumber, tomato, eggplant, and chili produce two virulence factors, pyocyanin, and siderophores (Ambreetha et al., 2021). Our current work extends our previous findings by characterizing the pathogenic phenotypes of those strains. Specifically, we assessed their ability to swarm, form biofilms, produce virulence factors, and kill other microbes and a select nematode.

### Vegetable-associated *P. aeruginosa* strains harbor multiple virulence traits

The *P. aeruginosa* strains tested in this study harbored an arsenal of virulence attributes. These include biofilm formation, swarming motility, rhamnolipid production, and lytic activity (hemolysis, proteolysis, and lipolysis).

#### Biofilm

Three endophytic (PPA03/cucumber, PPA08/tomato, PPA10/tomato), and two rhizospheric strains (PPA13/eggplant, and PPA14/eggplant) produced high levels of biofilm (Fig. 1). In agricultural plants, such as soybean, mung bean, sorghum, and tomato, biofilm-forming *P. aeruginosa* alleviates abiotic stress and enhances plant growth (Ali et al., 2009; Tank and Saraf, 2010; Sarma and Saikia, 2014; Kumawat et al., 2019). In the clinical setting, biofilm-forming *P. aeruginosa* is a dreaded pathogen and accounts for significant mortality in patients with critical pulmonary conditions (Römling et al., 1994; Singh et al., 2000; Nixon et al., 2001; Bjarnsholt et al., 2009). This is the first report to show that the endophytic *P. aeruginosa* strains present in cucumber (PPA03) and tomato (PPA08) can form biofilms comparable to clinical strains (Fig.1).

#### Swarming motility

There are no previous reports on the ability of plant-associated *P. aeruginosa* strains to swarm. In this study, four endophytic *P. aeruginosa* strains (PPA08, PPA10/tomato; PPA16, PPA18/chili) showed extensive swarming (Fig. 2). The tendril tip of the swarming bacteria possesses mobile cells that can quickly spread over any surface (Tremblay and Déziel, 2010). In the murine model system, it has been demonstrated that pathogenic *P. aeruginosa* swarms to disseminate in the host (Coleman et al., 2020a). Previous reports on clinical strains suggested that swarming motility might be associated with the expression of virulence factors (Overhage et al., 2008; Coleman et al., 2020b). In our study, the four superior swarmers exhibited lytic activity (α-hemolysis, proteolysis, and lipolysis) and comparatively higher antagonism against phytopathogens (Fig. 2, 4; Table 2).

#### Lytic activity

There were no previous reports on the hemolytic, proteolytic, or lipolytic capability of plant-associated *P. aeruginosa*. In this study, 10 of the 18 plant-associated strains exhibited α-hemolytic activity (Table 2). Hemolysin is an extracellular toxin produced by pathogenic bacteria to lyse host erythrocytes thereby facilitating tissue invasion (Goebel et al., 1988). Previous studies have demonstrated that in human infection, *P. aeruginosa* releases hemolysins to alter host lung physiology. This in part accounts for the serious morbidity and mortality associated with this bacterium (Darby et al., 1999; Wargo et al., 2011). In addition, *P. aeruginosa* extracellular lipase and protease disrupt cell membrane integrity and inactivate immune components (Heck et al., 1986; Parmely et al., 1990; König et al., 1996; Barker et al., 2004; Pinna et al., 2008). In our current study, 13 *P. aeruginosa* PPA strains exhibited protease activity; four of which also had lipase activity (Table 2).

#### Rhamnolipid

Both the clinical and agricultural strains of *P. aeruginosa* studied released rhamnolipid (Fig. 3). Previous clinical studies have suggested that *P. aeruginosa* rhamnolipids alter the respiratory epithelium facilitating lung infiltration (McClure and Schiller, 1996; Zulianello et al., 2006). However, in the agricultural ecosystem, rhamnolipids produced from *P. aeruginosa* protect the host plant against fungal pathogens (Oomycetes, Ascomycota, and Zygomycetes) and green peach aphid (Kim et al., 2000; Kim et al., 2011; Yan et al., 2015; Sancheti and Ju, 2019). In the current study, we have observed that the clinical strains PAO1 and ATCC10145 are the superior rhamnolipid producers (Fig, 3B). ATCC9027 has been previously reported as a low rhamnolipid producer (Grosso-Becerra et al., 2016). In this study, 50% of the PPA strains had higher levels when compared to ATCC9027.

### *P. aeruginosa* exhibits antagonism against phytopathogens

*Pythium aphanidermatum, R. solani*, and *F. oxysporum* are globally distributed fungal pathogens that cause rotting, blight, and wilt, respectively, in many plant species (Parmeter, 1970; Martin and Loper, 1999; Michielse and Rep, 2009; Lodhi et al., 2013). *X. oryzae* is a devastating rice pathogen that causes bacterial leaf blight (Swings et al., 1990). Previous reports have described that *P. aeruginosa* in agricultural ecosystems indirectly contributes to plant growth by inhibiting these harmful pathogens (Ali Siddiqui and Ehteshamul-Haque, 2001; Yasmin et al., 2014; Durairaj et al., 2017). The three control isolates of human origin, PAO1, ATCC10145, and ATCC9027, have never previously been tested for their ability to inhibit phytopathogens. Our current work demonstrates that both clinical and agricultural *P. aeruginosa* strains antagonize the tested fungal and bacterial phytopathogens (Fig. 4 and 5). This is unsurprising considering the number of virulence factors harbored by these strains (Fig. 1 to 3; Table 2). The secondary metabolites, pyocyanin and rhamnolipid, are implicated as the major determinants of *P. aeruginosa* antagonism (Kim et al., 2011; Sudhakar et al., 2015; Mahmoud et al., 2016; Chen et al., 2017; DeBritto et al., 2020). The strains tested in this study produced both pyocyanin and rhamnolipids (Fig. 3; Ambreetha et al., 2021) which might have contributed to anti-microbial virulence (Fig.4 and 5). Compared to PAO1, nearly 90% of the plant-associated strains had higher antagonism against the bacterial pathogen (Fig. 4D). In fungal system, the clinical strains had significantly higher virulence than most of the plant-associated strains (Fig. 4A to C). We suggest using the phytopathogenic fungi as a simple eukaryotic model system to test *P. aeruginosa* pathogenicity.

### Vegetable-associated *P. aeruginosa* induces mortality in *C. elegans*

The pathogenicity of *P. aeruginosa* in mammals is often assessed based on its lethality against *C. elegans* (Mahajan-Miklos et al., 1999; Tan et al., 1999). Virulent strains of *P. aeruginosa* accumulate in the nematode’s gut and slowly cause death (Tan et al., 1999; Kirienko et al., 2014). However, the non-pathogenic bacteria do not hinder the growth and development of *C. elegans* (Andrew and Nicholas, 1976). In this investigation, nematode mortality caused by the agricultural strains was considerably lower when compared to the clinical isolates. The most virulent agricultural strains (PPA03/cucumber; PPA08, and PPA10/tomato; PPA13, and PPA14/eggplant; PPA16, and PPA18/chili) induced mortality in 30-40% of the nematode population (Fig. 6). Despite the multiple virulence factors observed in the plant-associated strains, the mortality of *C. elegans* was higher (50-100%) when fed with clinical strains. The reduced virulence of the plant-associated *P. aeruginosa* strains suggests that the clinical isolates might have evolved to be more pathogenic to survive within the eukaryotic system. Pathoadaptive assays would reveal if these plant-associated strains can evolve into a more pathogenic form under the right conditions.

## Conclusion

To date, many studies have characterized the destructive virulence factors of human-associated, animal-associated, and environmental *P. aeruginosa* strains (Jaffar-Bandjee et al., 1995; Alonso et al., 1999; Vives-Flórez and Garnica, 2006; Zulianello et al., 2006; Balasubramanian et al., 2012; Hall et al., 2016; Moradali et al., 2017; Ruiz-Roldán et al., 2020). However, limited studies have demonstrated the ability of agricultural *P. aeruginosa* strains to infect animals and humans (Lebeda et al., 1984; Kumar et al., 2013). In this investigation, we have shown the presence of extremely virulent and lowly virulent *P. aeruginosa* strains in the rhizospheric and endophytic niches of four vegetables (cucumber, tomato, eggplant, and chili). Virulence was not correlated with the respective niche. The less virulent strains may be long-time soil dwellers and the extensively virulent strains might be human- or animal-adapted ones that got recently introduced into the agricultural ecosystem. These virulent strains may have entered the agricultural ecosystem through animal excreta or irrigation water with run-offs from nearby sewage systems (Wheater et al., 1980; Mavrodi et al., 2012; Slekovec et al., 2012; Orlofsky et al., 2016). Comparative genomic analyses will reveal the molecular adaptations contributing to the variation(s) among the agricultural strains. In the future, the pathoadaptive ability of the avirulent strains should be tested to find out if they could evolve into pathogens under selective conditions. Overall, this study reveals that agricultural plants harvested directly from soil could be a potential source for transmission of *P. aeruginosa* to humans. Farmworkers and consumers face risk of *P. aeruginosa* related infections, which are lethal in vulnerable individuals. To the best of our knowledge, this study is the first comprehensive attempt to show that *P. aeruginosa* strains residing within the internal tissues and rhizosphere of edible vegetables harbor multiple virulence factors critical for human infection.

## Experimental Procedures

### Bacterial strains and culture conditions

Plant-associated *P. aeruginosa* strains isolated and characterized in our previous study were used as test strains (Ambreetha et al., 2021). Clinical strains of *P. aeruginosa*, PAO1, ATCC10145, and ATCC9027 were used as controls (Table 1). All *P. aeruginosa* strains were periodically sub-cultured and grown in Pseudomonas agar (for pyocyanin) medium (PAP, Himedia) at 37°C. A plant pathogenic bacterium*, Xanthomonas oryzae,* was cultured in a nutrient agar medium at 37°C. Plant pathogenic fungi, *Pythium aphanidermatum, Rhizoctonia solani, and Fusarium oxysporum* were cultured in potato dextrose agar medium at 37°C.

### Nematode strain and culture conditions

*Caenorhabditis elegans* N2 hermaphrodite strain was used in this study (Brenner, 1974). The worms were periodically cultured in nematode growth medium (NGM), overlaid with *Escherichia coli* strain OP50, and maintained at 20°C (Brenner, 1974).

### Biofilm production

*In vitro* biofilm production by the *P. aeruginosa* strains was quantified using microtiter assay (O’Toole, 2011). Overnight cultures of the *P. aeruginosa* strains (25 µl, OD660∼0.5) were inoculated into microtitre wells containing 225 μl of LB broth. Three sets of microtitre plates were inoculated and incubated for three different time intervals (24, 48, and 72 hours). After the incubation period, planktonic cells were transferred to a new microtitre plate and A660 was measured (Spectramax^®^ i3x, USA). Biofilms stuck to the plates were washed twice with sterile H2O and flushed with 0.1 % of crystal violet. The plates were incubated for 10-15 minutes at room temperature and gently washed twice with sterile H2O. The stained plates were allowed to dry overnight at room temperature. The next day, 30% acetic acid was added to the well to dissolve the biofilms, and absorbance was measured @ 550 nm. The ratio between the biofilm and planktonic populations was determined at three time points (24, 48, and 72 h; O’Toole, 2011). The experiment was repeated thrice and the results were represented as the biofilm to planktonic ratio.

### Swarming motility

Swarming motility of the *P. aeruginosa* strains was assessed by adding 10 µl of 24 h-old test strains (OD660 ∼ 0.5) on modified M9 plates with 0.5% agar (Tremblay and Déziel, 2008). The diameter of the bacterial tendrils extended on the plates due to swarming was measured, and the percentage of plates swarmed within 48 h of incubation was estimated (Tremblay and Déziel, 2008). The experiment was repeated thrice and three different diameters were measured every time. Results were represented as the percentage of the plate area swarmed in 48 h.

### Rhamnolipid

#### Qualitative assay

A CTAB agar test was done to qualitatively assess the *P. aeruginosa* strains for rhamnolipid production (Siegmund and Wagner, 1991). In brief, the culture supernatants of the test strains were filtered using 0.45 µm filters. Ten microliters of the cell-free extracts were added to 0.2 cm wells on CTAB-methylene blue agar plates and incubated at 37°C for 24 h. If rhamnolipid (anionic surfactant) was present in the supernatant, it reacted with the CTAB (cationic surfactant), resulting in an insoluble complex. The strains were scored positive based on the formation of a dark blue precipitated zone around the culture wells. The experiment was repeated thrice to confirm result consistency.

#### Quantitative assay

The strains were grown in phosphate limited protease peptone ammonium salts (PPGAS) broth supplemented with 2% (v/v) sunflower oil at 37°C to induce rhamnolipid production for seven days (Zhang and Miller, 1992). We used the chloroform-methanol extraction method for rhamnolipid separation (Zhang and Miller, 1992). In brief, the cell-free culture supernatant was acidified to pH2 with 12 M hydrochloric acid. The lipids were extracted using a chloroform-methanol (2:1) mixture and concentrated through evaporation. Concentrated rhamnolipids were gravimetrically quantified (Gunther et al., 2005). The experiment was repeated thrice and results were presented as µg/ml of the culture supernatant.

### Lytic activity

#### Hemolysis

The ability of the *P. aeruginosa* strains to lyse blood cells was assessed by streaking the overnight cultures (OD660 ∼ 0.5) on nutrient agar plates containing 5% sheep blood (Williams and Harper, 1947). The plates were incubated for 24 h at 37°C. Green discoloration of the blood with a mild halo zone was noted as α-hemolysis, and the absence of lytic activity was noted as γ-hemolysis. The experiment was repeated thrice for consistency.

#### Lipolysis and proteolysis

The lipolytic activity of the *P. aeruginosa* strains was assayed using 1% tributyrin as a substrate (Atlas, 1993; Georgescu et al., 2016). Strains were considered positive for lipolytic activity if an opaque precipitate formed around the bacterial colonies. The proteolytic behavior of the strains was assayed using 3% skim milk and 3% gelatin (1;1 ratio) as the substrates (caseinase, and gelatinase activity, respectively). Formation of halo zones around the colonies were indicative of casein proteolysis and gelatin hydrolysis (Atlas, 1993; Georgescu et al., 2016). The strains were scored based on the intensity of lysis (mild lysis, heavy lysis, or no lysis). The experiment was repeated thrice for consistency.

### Antagonistic activity

#### Antifungal antagonism

The antagonistic potential of the *P. aeruginosa* strains against three phytopathogenic fungi (*Pythium aphanidermatum, R. solani*, and *F. oxysporum*) was assessed by dual-culture assay (Sakthivel and Gnanamanickam, 1986). In brief, fungal discs were placed on one corner of potato dextrose agar medium in 90 mm Petri plates. *P. aeruginosa* strains were streaked 3 cm away from the fungal disc. Plates were incubated @ 37°C for seven days. Inhibition in mycelial growth, as influenced by the *P. aeruginosa* strains, was recorded. The percentage inhibition was estimated based on the standard formula, 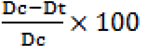, where Dc is the diameter of the fungal mycelium in the control plate and Dt is the diameter of the fungal mycelium as influenced by the test strains (Riungu et al., 2008). The experiment was repeated thrice for consistency.

#### Antibacterial antagonism

Antibacterial effect of the test strains against *Xanthomonas oryzae* pv. *oryzae* was estimated by cross streak assay (Lertcanawanichakul and Sawangnop, 2011). In brief, *P. aeruginosa* strains were streaked at the center of nutrient agar plates and incubated for 24 hrs. After 24 hours, *X. oryzae* was streaked perpendicular to the central streak and the plates were incubated for another 24 hrs @37°C. Inability of the target pathogen to grow in the confluence area was recorded after incubation. The percentage inhibition of *X. oryzae,* as influenced by the *P. aeruginosa* strains, was calculated based on the standard formula, 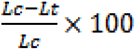, where Lc is the length of the *X. oryzae* grown in the control plate and Lt is the length of the *X. oryzae* as influenced by the test strains (Lo Giudice et al., 2007). The experiment was repeated thrice for consistency.

### *C. elegans* killing assay

The ability of the *P. aeruginosa* strains to induce death in *C. elegans* was demonstrated via a slow-killing assay (Tan et al., 1999). *C. elegans* gravid adults were treated with 1N NaOH and 5% sodium hypochlorite (1:1) solution (Brenner, 1974). The eggs were allowed to hatch in M9 buffer, and 24 h later the emerged L1-worms were released over a lawn of *E. coli* OP50 (Brenner, 1974; Adonizio et al., 2008). These synchronized L1- worms were grown up to the L4-stage. We prepared slow-killing plates with NGM (Tan et al., 1999) seeded with overnight cultures (OD660 ∼ 0.5) of OP5O, PPA strains, and clinical strains of *P. aeruginosa.* The plates were incubated at 37°C for 24 h. The L4- worms (20 per plate) were introduced into these plates and incubated at 20°C (Brenner, 1974). The viability of the nematodes, as influenced by the tested bacterial strains, was recorded every 24 h for five consecutive days. Worms that did not respond to physical stimuli were scored as dead. The death of worms on the OP50 plate was scored as natural mortality (negative control). The experiment was repeated thrice for consistency.

### Statistics and reproducibility

All experiments were performed in triplicates. All data were subjected to a one-way analysis of variance (ANOVA) with a P-value of 0.05, and Duncan’s multiple range test was performed between individual means to reveal any significant difference (XLSTAT, version 2010.5.05 add-in with Windows Excel). Principal coordinate analysis (PCoA) based on Euclidean distance was carried out using NCSS 2020 statistical software (NCSS, Kaysville, USA) to cluster the *P. aeruginosa* strains based on their antagonism against phytopathogens. Data analysis and scientific graphing were done in OriginPro version 8.5 (OriginLab®, USA).

## Acknowledgment

No grant supported this work. SA was partially funded by the Fulbright Doctoral Nehru Research Fellowship by the U.S Department of State’s Bureau of Educational and Cultural Affairs and United-States India Educational Foundation (ID. PS00299273). SA also received Science and Engineering Research Board-International Travel Support funded by the Department of Science and Technology, India (No: ITS_2019_002449). We thank Dr. Sriyutha Murthy (Indira Gandhi Centre for Atomic Research, Kalpakkam, Tamilnadu, India) for providing *P. aeruginosa* strain PAO1 and Dr. Kavitha Babu (Department of Biological Sciences, Indian Institute of Science Education and Research, Mohali) for providing the *C. elegans* nematode.

## Author Contributions

The experiments were conceived and designed by SA and DB. The samples were processed by SA and PM. The experiments were performed by SA. Critical analyses of the data were done by SA, KM, and DB. The manuscript was prepared by SA and KM. Finally, all authors were involved in the critical review of this paper.

### Ethical Approval

There were no human or animal subjects involved in this study.

## Data Availability

All sequence data generated in this study were deposited in NCBI GenBank (Accession no. MT734694 to MT734711).

